# The Fat4 intracellular domain controls turnover of Fat4/Dchs1 planar polarity membrane complexes

**DOI:** 10.1101/2024.05.25.595882

**Authors:** Yathreb Easa, Olga Loza, Roie cohen, David Sprinzak

## Abstract

The Fat/Dachsous (Ft/Ds) pathway is a highly conserved pathway regulating planar cell polarity (PCP) across different animal species. Proteins from the Ft and Ds family are large transmembrane protocadherins that form heterophilic complexes on the boundaries between cells. Fat4 and Dchs1, the main mammalian homologues of this pathway, have been implicated in PCP in various epithelial tissues and were shown to form extremely stable complexes at the boundaries between cells. It is unclear however, what are the dynamics controlling such stable boundary complexes, and how the formation and turnover of these complexes is regulated. Here, we use quantitative live imaging to elucidate the role of the intracellular domains (ICD) of Fat4 and Dchs1 in regulating Fat4/Dchs1 complex dynamics. We show that removing the ICD of Fat4 results in a reduction of both Trans-endocytosis (TEC) of Dchs1 into the Fat4 cells and boundary accumulation, but does not affect the diffusion of the complexes at the boundary. We further show that the ICD of Fat4 controls the turnover rate of Fat4/Dchs1 complexes. Finally, we find that while actin polymerization is required for maintaining the boundary accumulation of Fat4/Dchs1 complexes, we do not identify correlations between Fat4/Dchs1 complexes and local actin accumulation. Overall, we suggest that the Fat4 ICD is important for the turnover and plasticity of the highly stable Fat4/Dchs1 complexes associated with PCP.

**Statement of Significance:** The purpose of this study is to elucidate the dynamics leading to the formation and maintenance of Fat4/Dchs1 complexes at the cell boundary and how it is affected by the ICD of Fat4 in mammals. The insights from this work are important for obtaining a mechanistic molecular framework for understanding formation of planar cell polarity as well as for highlighting how cell boundary membrane complexes are regulated.

## Introduction

Planar cell polarity (PCP) is the process by which sheets of cells, typically in the form of epithelial cell layers, acquire an orientation within the plain of the sheet (1). Coordinated polarity generates an organized tissue geometry required for the proper function of many developing organs (2). The polarity of epithelial cells is usually represented by orientation of external structures, such as trichomes and bristles in *Drosophila*, sensory cells in the inner ear and skin hairs in vertebrates (3–5). These types of PCP are regulated by transmembrane protein complexes which belong to two different pathways, the core pathway (Frizzled/Van-Gogh pathway) and the Fat/Dachsous (Ft/Ds) pathway. Both were originally discovered in *Drosophila* but are evolutionarily conserved in higher vertebrates (2, 6–7).

In *Drosophila*, the Ft/Ds pathway components include the large atypical protocadherins Ft and Ds (8). Ft and Ds are transmembrane proteins that bind to each other at the cell surface and take part in heterophilic interactions resulting in trans-hetero-complexes on the boundary between cells (9). Ft/Ds complexes are asymmetrically distributed on the boundaries between cells (localized in one direction (2)). This asymmetry is associated with determining the direction of PCP.

The mammalian homologues of Ft and Ds include Fat1-Fat4 and Dchs1-Dchs2. However, Fat4 and Dchs1 are the most studied members in mammals, consistent with their high degree of homology to *Drosophila* Ft and Ds. Fat4 and Dchs1 are also the most widely expressed, and have the strongest knockout phenotypes (10). Fat4 and Dchs1 null mice show complex morphological abnormalities in the kidney, inner ear, bone, brain, lymph node, vertebrae, skeleton and more (11). In humans, mutations in Fat4 and Dchs1 were recently linked to various cancers and abnormal brain development (12).

In *Drosophila*, the Ft ICD has been shown to be essential for both growth control and PCP (13–15). In fact, different domains within Ft ICD have been associated with either growth control function or PCP (14,15). Replacing the ICD of Ft with that of the mammalian Fat4 rescued PCP activity in *Drosophila* but not growth control phenotypes, suggesting a role for Fat4 ICD on PCP. However, how the Fat4 ICD regulates Fat4/Dchs1 boundary complexes and PCP in mammals is not understood.

We have previously used mammalian cell culture system to show that Fat4 and Dchs1 form stable membrane complexes on the boundary between cells, and suggested that the formation of these stable boundary complexes is required for PCP. We have further shown that Dchs1 trans-endocytose into the Fat4 cells, while no TEC of Fat4 is observed into the Dchs1 cells (16).

Given the significant gaps in our understanding of how the Fat4/Dchs1 pathway regulates PCP in mammals, we focus here on determining the role of the ICDs of Fat4 and Dchs1 during the formation of boundary complexes in mammals. We use a cell culture assay to study how the ICD of Fat4 and Dchs1 regulate boundary accumulation, stability, and turnover. We show that boundary accumulation still occurs without the ICD of either Fat4 or Dchs1, but TEC is significantly reduced when Fat4 lacks its ICD. We further show that Fat4/Dchs1 complexes are stable on the boundary between cells and that removing the ICD of Fat4 does not affect the diffusion of the complexes at the boundary. Additionally, our results suggest that the turnover rate of Fat4/Dchs1 complexes are faster with wildtype (WT) Fat4 compared to Fat4 lacking ICD. Finally, we show that while actin polymerization is required for maintaining the boundary, we do not see enhancement of actin dynamics next to the boundary where accumulation is observed, regardless of whether Fat4 ICD is present or not.

## Methods

### Cells and Plasmids

Human embryonic kidney 293T cells (HEK293T-ATCC CRL3216) were grown in adherent cultures in Dulbecco’s Modified Eagle’s Medium (DMEM) supplemented with 10% FBS. All cells were cultured in a humidified atmosphere of 5% CO2 at 37°C. Stable and transient transfections were performed using TransIT-LT1 reagent (Mirus Bio, Madison, Indiana) or Lipofectamine 3000 (Thermo Fisher Scientific, Waltham, MA USA) according to the manufacturer’s instructions.

Transfection was performed with 1ug of Fat4 WT, Dchs1 WT, Fat4 ΔICD and Dchs1 ΔICD plasmids. For the mCerulean3-LifeAct plasmid we used a mixture of 200 ng of the desired plasmid with 800 ng of an empty pORI vector (17). Two days after transfection cell were transferred to a 6-well plate and placed under selection of 100 µg/ml Zeocin (InvivoGen, San Diego, USA) for Fat4-Citrine constructs, 5 µg/ml Blasticidin and 100 µg/ml Hygromicin (AG Scientific, San Diego, USA) for Ds1-mcherry constructs, 100 µg/ml Zeocin and 5 µg/ml Blasticidin for the mCerulean3-LifeAct constructs.

To create single cell colonies after selection was applied, the cells were diluted to a concentration of 2 cells/ml and transferred into a 96-well plate. After a period of two weeks, the plates were screened for positive clones, which were transferred to a new plate for further growth. For the double stable cell lines of Fat4-WT/mCerulaen3 - LifeAct and Fat4-ΔICD/mCerulean3-LifeAct were used as mixed populations.

All plasmids were constructed using standard cloning techniques. Fat4-Citrine-WT, Dchs1-mCherry-WT are based on constructs developed in (16). Fat4 ΔICD Citrine, Dchs1 ΔICD mCherry contain the full extracellular and transmembrane domains but only the first nine amino acids of the intracellular domains of Fat4 and Dchs1. respectively. The mCerulean3-LifeAct was ordered from addgene (plasmid number - 54721). The selection marker G418 was replaced by Blasticidin, then co-transfected with Fat4-Citrine-WT and Fat4-ΔICD-Citrine. All plasmids used in this study are listed in Supplementary Table 1 (also contains links to full sequences).

### Co-culture assays

For the co-culture TEC assay, boundary accumulation, FRAP analysis, mCerulean3-LifeAct, Lat-A and Ca^+2^experiments, were seeded in glass bottom 24-well plates (Cellvis, California, USA) with 1:1 ratio between Fat4 and Dchs1 expressing cells. The plates were covered with Poly-L-lysine solution (10 % (v/v) in DDW H2O) (Sigma Aldrich, Saint Louis, USA) to improve cell adherence. Cells were grown for 24 hours and directly prior to imaging the media was replaced with low fluorescence imaging media (αMEM without Phenol red, ribonucleosides, deoxyribonucleosides, folic acid, biotin and vitamin B12, Biological Industries, Israel). For the Lat-A experiments Lat-A (abcam-ab144290, Cambridge, MA,USA) were added for different periods of time. For the Ca^+2^ washout experiments Cells were grown for 24 hours and then the growth medium was replaced to medium without Ca^+2^ (DMEM-Gibco) for a different periods of time. After that, the cells were washed with PBS and fixed with 4% paraformaldehyde in PBS for 10 minutes at room temperature.

### Flow Cytometry

Cells were seeded in 6 well plates at approximately 70% confluence 24 hours before FACS. Directly prior to FACS, cells were trypsinized, spun at 1000 rpm for 5 min, and resuspended in 200 µL of FACS buffer. FACS buffer consisted of PBS with 1% FCS serum, and 5mM EDTA. Flow cytometry was performed using a Cytoflex5L flow cytometer (Beckman Coulter). Kaluza software was used to analyze the data.

### Imaging of cells

Cells in co-culture experiments were imaged using a Zeiss LSM 880 confocal microscope using a 514nm laser for Citrine, a 561nm laser for mCherry and a 405nm laser for mCerulean3-LifeAct. The images were taken using a Plan-apochromat 63× oil-immersion objective with a numerical aperture (NA) of 1.4. Microscope was equipped with a 37°C temperature–controlled chamber and a CO2 regulator providing 5% CO2.

### FRAP experiments on the accumulating boundaries

FRAP experiments on the accumulating boundaries were performed using Andor revolution spinning disk confocal microscope with DPSS CW 515nm (50% power) and 561nm (50%power) 50mW lasers (Andor, Belfast, Northern Ireland). Andor revolution spinning disk confocal microscope supplied with FRAPPA device (Andor, Belfast, Northern Ireland). Photobleaching was performed with 100% power of the 445nm laser for a total bleach time of 150ms (3 repeats of 50ms). One image was taken before the bleach and 50-90 images were taken after the bleach every 10 seconds.

### TEC analysis

To estimate the amount of TEC in co culture cells of Fat4 and Dchs1 expressing cells, we calculated the amount of Dchs1 inside Fat4 expressing cells. As Fat4 is marked by Citrine and Dchs1 by mCherry, we assumed that the level of TEC is proportional to the total signal of the red channel inside the Fat4 Citrine expressing cells. To estimate that, we used intensity thresholding in the green channel to create a binary mask representing the Fat4 cells region. The estimated TEC per cell was defined as the total intensity of the red channel inside the masked region, divided by the number of cells.

### Boundary accumulation analysis

To estimate the colocalization of Fat4 and Dchs1 on the boundaries between the cells, we first manually traced the boundaries where accumulation is observed using a custom-made MATLAB code. To create a binary mask that represents the boundary regions, we dilated the manually delineated lines by a radius that is slightly larger than the characteristic boundary width. This edge mask was used to separately estimate the amount of Fat4-Citrine and Dchs1-mCherry on the boundaries. The boundary accumulation was then calculated by multiplying both channels by the edge mask and summing the red and green pixel intensity. To estimate the colocalization on the boundaries we first applied gaussian blurring on both channels (intended to close the gap between the centers of the boundaries between two channels). The gaussian width (sigma) was chosen to be the distance between the centers of the boundaries. The blurred channels were multiplied by each other, then by the edge mask, and then summed and divided by the total length of the boundaries.

The MATLAB code used to analyze the TEC and boundary accumulation experiments can be found at https://github.com/Roie-Cohen/yathrebe_tec_analysis

## Results

### Characterizing the role of the intracellular domain of Fat4 and Dchs1

Given the identified role of the ICD of *Drosophila* Ft and Ds, we wanted to test its role in the mammalian homologs, Fat4 and Dchs1. We have previously developed a cell culture assay for studying the interactions between Fat4 and Dchs1 (16,18). This assay is based on HEK293T cells expressing Fat4-Citrine, or Dchs1-mCherry placed under a constitutive expression promoter (CMV). We performed co-culture experiments of Fat4-Citrine-WT: Dchs1-mCherry-WT and follow the interactions in a confocal microscope (Fig. 1A, Supplemental Fig. S1). Such co-culture experiments showed a strong accumulation of Fat4-Citrine and Ds1-mCherry at heterotypic junctions, as previously observed (yellow arrow in Supplemental Fig. S1A). These images also showed TEC of Dchs1-mCherry into the Fat4-Citrine cells as evident from numerous vesicles containing Dchs1-mCherry in the Fat4-Citrine cells (white arrow in Supplemental Fig. S1). Almost no trans-endocytosis in the reverse direction, from Fat4-Citrine into the Dchs1-mCherry, was observed. These trans-endocytosed vesicles show similar dynamics to endocytic vesicles when live imaging is performed (white arrows in Supplemental Fig S1B).

**Figure 1:**
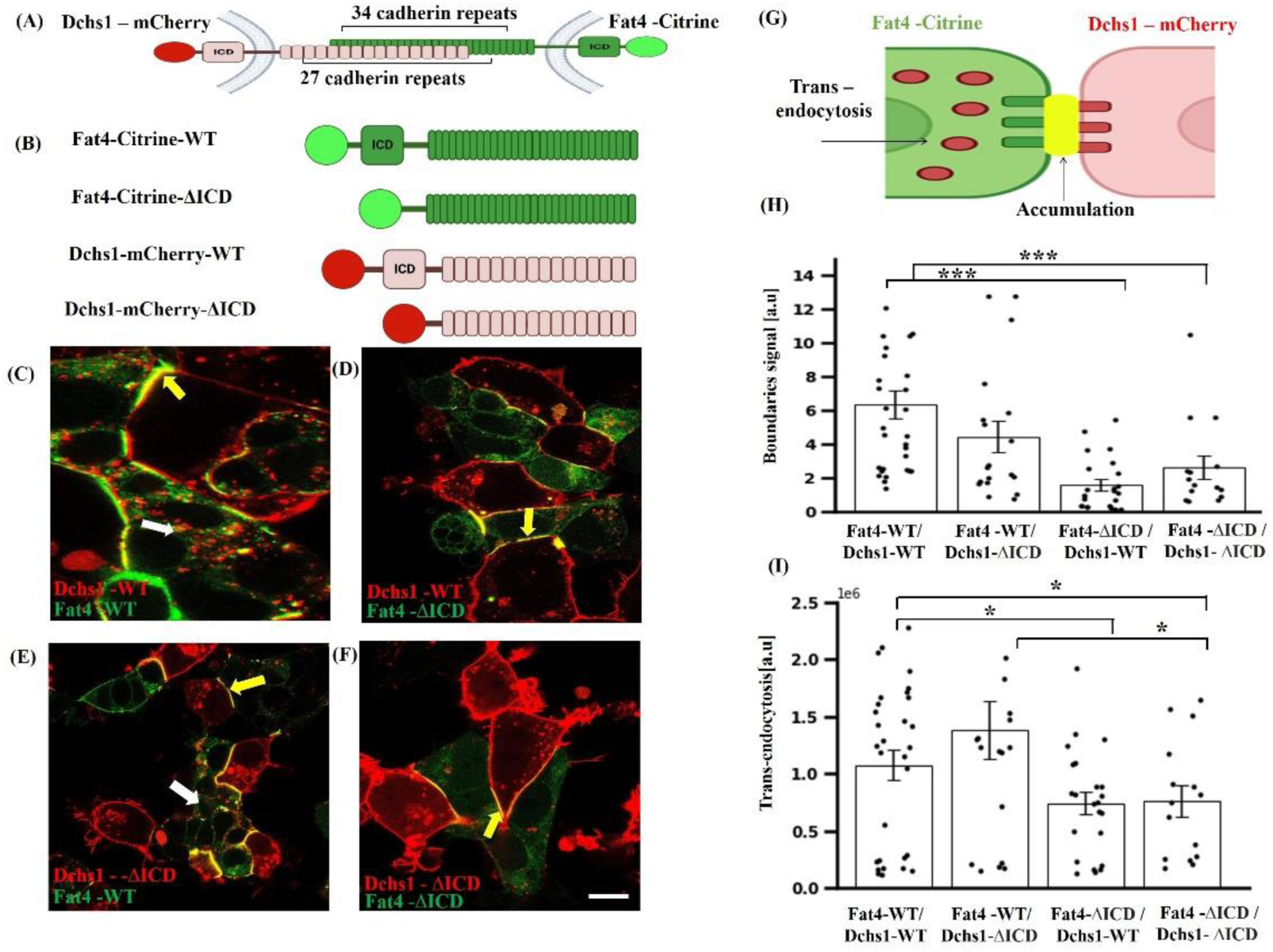
The ICD of Fat4 affects accumulation and trans-endocytosis of Fat4/Dchs1 boundary complexes. **(A)** Schematic illustration of a Fat4/Dchs1 boundary complex. **(B)** Schematic of the Fat4-Citrine-WT, Fat4-Citrine- ΔICD, Dchs1-mCherry-WT, and Dchs1-mCherry-ΔICD constructs. **(C-F)** A co-culture of Fat4-Citrine-WT or Fat4- Citrine-ΔICD cells (green) with Dchs1-mCherry-WT or Dchs1-mCherry-ΔICD cells (red). All co-culture combinations exhibit boundary accumulation (yellow arrows). Trans-endocytosis (TEC) is predominantly observed in co-cultures with Fat4-ΔICD. Co-cultures with Fat4-WT and Dchs1-ΔICD exhibits TEC levels similar to those of WT-WT cultures. **(G)** Schematic illustration showing the boundary accumulation and TEC measured in co-cultures of Fat4-Citrine (green) and Dchs1-mCherry (red). Yellow boundary represents accumulation in heterotypic boundaries, vesicles represent trans-endocytosis of Dchs1-mCherry into the Fat4-Citrine cells. **(H-I)** Quantitative analysis of the boundary complex formation (H, multiplication of Dchs1-mCherry and Fat4-Citrine signals on the boundary) and TEC (I, showing Dchs1-mCherry fluorescence in Fat4-Citrine cells) in the different co-cultures. See methods for how both quantities are defined. Data points show mean values from n=30 for Fat4-Citrine-WT:Dchs1-mCherry-WT, n=23 for Fat4-Citrine-ΔICD:Dchs1-mCherry-WT, n=19 for Fat4-Citrine-WT:Dchs1-mCherry-ΔICD and n=15 for Fat4-Citrine-ΔICD: Dchs1-mCherry-ΔICD from three independent experiments. Error bars represent S.E.M. *P < 0.05; ***P < 0.001. Scale bars-10 μm.

### The ICD of Fat4, but not of Dchs1, is required for TEC

To test the role of the intracellular domains of Fat4 and Dchs1 on boundary dynamics and TEC, we generated new constructs that lack the ICD of Fat4 and Dchs1 (Fig. 1B). We generated stable cell lines expressing either the wildtype (WT) constructs or the ones lacking the ICD (ΔICD). We then performed a comparative imaging assay looking at both boundary accumulation and TEC.

We have observed that boundary accumulation of Fat4 and Dchs1 did occur when we co-cultured WT constructs with constructs lacking their ICD, indicating that the ICD of Fat 4 and Dchs1 is not required for boundary accumulation (yellow arrows in Fig. 1C-F). To test if the level of boundary accumulation was different between Fat4 and Dchs1 constructs with or without their ICD we have developed a quantitative analysis that used a semi-automated analysis of boundary accumulation. In this assay, we quantitatively measured the green and red fluorescence along the boundaries showing accumulation, and used the multiplication of both signals as a measure for accumulation (see methods and Fig.1G).

The analysis revealed a significant reduction in the boundary fluorescence when WT-Dchs1 was co-cultured with Fat4-ΔICD instead of Fat4-WT (Fig. 1D, H, Supplemental Fig. S2). We do observe a somewhat weaker reduction in boundary accumulation when the Dchs1 ICD is missing (Fig. 1E, H). The effect of removing both ICDs is similar to that of removing only the Fat4-ΔICD (Fig. 1F, H). These results suggest that the ICD of Fat4 (but likely not the ICD of Dchs1) is important for achieving stronger accumulation of Fat4/Dchs1 complexes at the boundary.

We next looked at the effect of the ICD of Fat4 and Dchs1 on the TEC of Dchs1. To quantify TEC, we measured the Dchs1-mCherry fluorescence inside the Fat4-cirine cells (see methods). The analysis revealed that there is a significant decrease of the TEC of Dchs1-mCherry into Fat4-Citrine cells when Fat4 ICD is removed (Fig. 1D, I). This reduction is observed also when the co-culture is performed with Dchs1-ΔICD (Fig 1F,I). The removal of the Dchs1 ICD alone does not affect TEC. These results suggest that the ICD of Fat4, but not of Dchs1, is required for the TEC process.

### The expression and subcellular localization of the Fat4 and Fat4-ΔICD are similar

A possible explanation for the different behaviors between Fat4-WT and Fat4-ΔICD could be that the different cell lines express the variants at different levels or that the subcellular localization is different between the two variants. To test the expression levels, we performed FACS analysis on the different cell lines we used (Supplemental Fig. S3). We found that there are no significant differences in the protein expression between Fat4 and Fat4-ΔICD. Similarly, there is no difference in expression between Dchs1 and Dchs1-ΔICD (Supplemental Fig. S3B). In addition, we have looked at the protein localization of the different variants of Fat4 and Dchs1 when not in co-culture. In general, the subcellular localization seems to be similar between the WT variants and the variants lacking the ICD (Supplemental Fig. S3A). These results suggest that the different behaviors are not related to the differences in the expression levels or in the subcellular localization.

### Fat4 and Dchs1 form stable complexes on the boundary even in the absence of the ICD of Fat4

We have previously shown that the formation of Fat4 and Dchs1 complexes significantly stabilizes these proteins on the cell membrane (e.g. reduce the diffusion coefficient), possibly through the formation of boundary clusters (16). We therefore decided to test the effect of the ICD of Fat4 on the stability of Fat4/Dchs1 complexes at the boundary between cells. In order to test stability of Fat4/Dchs1 complexes, we used fluorescence recovery after photobleaching (FRAP) experiments in regions where boundary accumulation was observed. In these assays we bleached boundaries where accumulation was observed using a blue 445nm laser, which successfully bleached the Fat4-Citrine fluorescence, but did not affect the Dchs1-mCherry, and tracked the recovery. We compared the recovery after photobleaching in Fat4-Citrine-WT: Dchs1-mCherry-WT boundaries (Fig. 2A-B, movie S1) to that in Fat4-Citrine-ΔICD: Dchs1-mCherry boundaries (Fig. 2D-E, movie S2). As controls, we also tracked the dynamics in boundaries that were not bleached to account for photobleaching due to imaging alone. We note that these controls without FRAP also exhibited significant photobleaching (Fig. 2C, F) hence the results of the FRAP experiments were always compared to these controls. (Fig. 2G).

**Figure 2:**
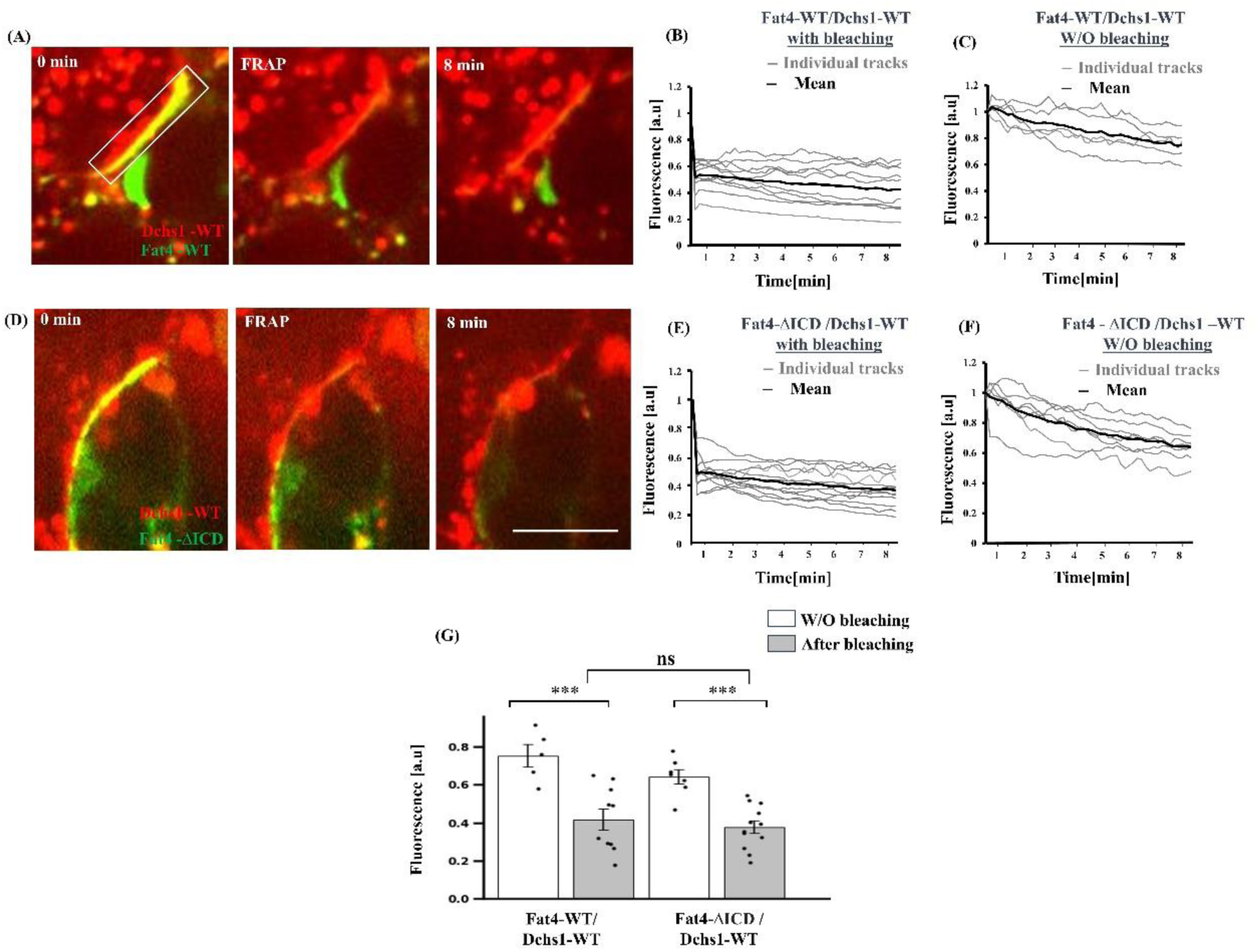
Fat4 ICD is not required for the stability of Fat4/Dchs1 complexes. **(A, D)** Filmstrips showing a fluorescence recovery after photobleaching (FRAP) experiment on a boundary exhibiting accumulation (yellow) of co culture of Fat4-Citrine-WT (green): Dchs1-mCherry-WT (red) (A) or Fat4-Citrine-ΔICD (green): Dchs1-mCherry-WT (red) (D). The region that was bleached is marked with white rectangle. **(B-C, E-F)** Graphs showing the fluorescence recovery as a function of the time (x-axis) in a co culture of Fat4-Citrine-WT: Dchs1-mCherry-WT (B) or Fat4-Citrine-ΔICD: Dchs1-mCherry-WT. (E) As a control we imaged boundaries without intentional photobleaching (C, F). The graphs with intentional photobleaching (B, E) do not exhibit any significant recovery, even after 8 minutes, regardless of the presence or absence of Fat4 ICD. **(G)** A graph showing the relative fluorescence after 8 minutes with photobleaching (green) or without intentional photobleaching (blue) in a co-culture of Fat4-Citrine-WT: Dchs1-mCherry-WT or Fat4-Citrine-ΔICD: Dchs1-mCherry-WT cells. Number of experiments: For (B-C), without photobleaching, n=5, with photobleaching n=10. For (E-F), without photobleaching, n=7, with photobleaching, n=12. Measured fluorescence was normalized by the pre-bleach fluorescence level. Gray lines: individual FRAP experiments. Black lines: average of all FRAP experiments. Error bars represent S.E.M.***P<0.001, ns indicates not significant p-value>0.05). Scale bars-10 μm.

Consistent with our previous results, our analysis demonstrates that Fat4 and Dchs1 form extremely stable complexes on the boundary, since no significant recovery is observed after bleaching of Fat4-Citrine-WT: Dchs1-mCherry-WT boundaries over time scales of minutes (Fig. 2B). Similarly, when the ICD of Fat4 was removed (tracking recovery in Fat4-ΔICD-Citrine: Dchs1-mCherry-WT) no significant recovery after bleaching was observed (Fig. 2E). These results showed that removing the ICD of Fat4 did not affect the stability of Fat4/Dchs1 complexes, and hence suggest that Fat4-ICD is not required for membrane stability of Fat4/Dchs1 complexes.

### The ICD of Fat4 is required for faster turnover upon depletion of Ca^+2^

Since Fat4 and Dchs1 are protocadherins, their binding is calcium (Ca^+2^) dependent (8,19). We therefore decided to test the effect of Ca^+2^ depletion on the turnover of Fat4/Dchs1 complexes and their accumulation on the boundary. To do that we performed Ca^+2^ switch on Fat4/Dchs1 co-cultures, by replacing the normal growth medium of the cells to medium without Ca^+2^ and tracking the number of boundaries exhibiting Fat4/Dchs1 accumulation per field of view. This was performed for both Fat4-Citrine-WT: Dchs1-mCherry-WT and in Fat4-Citrine-ΔICD: Dchs1-mCherry-WT boundaries (Fig. 3A). The analysis showed a significant reduction of the fraction of Fat4/Dchs1 boundaries already half an hour after depleting Ca^+2^ from the medium for both co-culture experiments. However, the reduction in the fraction boundaries in the experiment with Fat4-WT was significantly faster than in the experiments with Fat4-ΔICD. These results suggested that the ICD of Fat4 regulate the turnover of Fat4/Dchs1 complexes (Fig. 3B).

**Figure 3:**
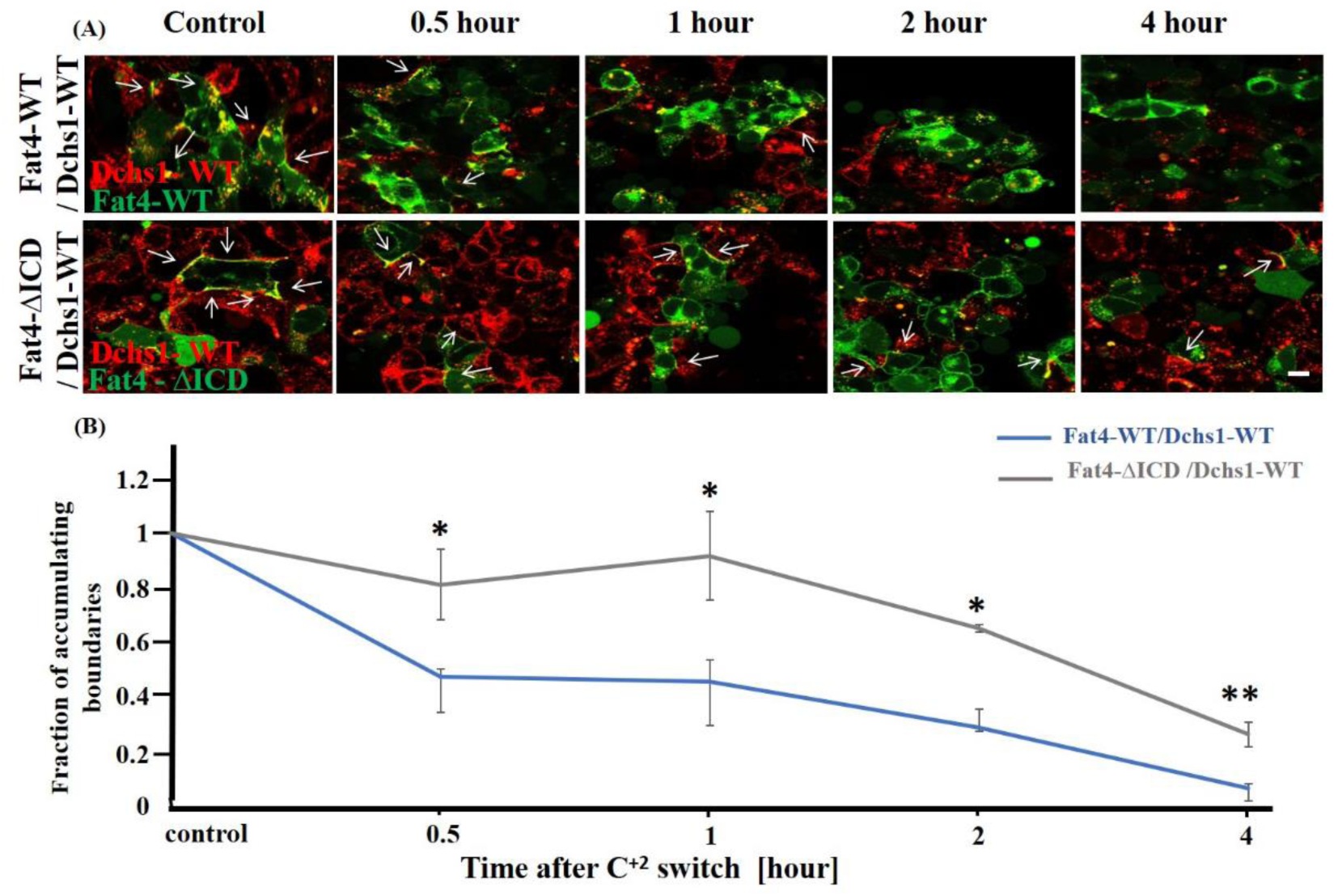
Fat4/Dchs1 turnover after Ca^+2^ depletion is slower in the absence of Fat4 ICD. **(A)**. Images from a co-culture of Fat4-Citrine-WT (green): Dchs1-mCherry-WT (red) cells (top) or Fat4-Citrine-ΔICD (green): Dchs1-mCherry-WT(red) cells (bottom) at different time points after switching to a medium without Ca^+2^. Control experiment represents the situation in normal growth medium. **(B)** A graph showing the fraction of accumulating boundaries (with respect to control) in co-culture of Fat4-Citrine-WT: Dchs1-mCherry-WT (blue), co-cultures of Fat4-Citrine-ΔICD: Dchs1-mCherry-WT (gray) at different time points after Ca^+2^ depletion. The fraction of accumulating boundaries normalized by the average number in the control. Number of images: for the first-row n=16,15,16,15,14, for the second-row n= 16,16,16,17,18, respectively. Images taken from three independent experiments. Error bars represent S.E.M.*P<0.05, **P.,0.01. Scale bars-10μm.

### Actin polymerization is required for maintaining the boundary

It has been shown that membrane complexes or adhesion molecules modulate adhesion through dynamic interactions with the actin cytoskeleton (19–20). To test if actin cytoskeleton is required for the formation or accumulation of Fat4/Dchs1 complexes, we used Latrunculin-A (Lat-A), a reagent that depolymerize actin filaments and is often used in experiments with live cells (22). We tested whether Lat-A affects the formation or accumulation of Fat4/Dchs1 boundaries in the presence or absence of Fat4 ΔICD (Fig. 4A). Our results revealed a significant reduction of the number of Fat4/Dchs1 boundaries both 0.5h and 1h after adding Lat-A in co-cultures of Fat4-Citrine-WT: Dchs1-mCherry-WT and Fat4-Citrine-ΔICD: Dchs1-mCherry-WT compared to the control (DMSO) (Fig. 4B-C). This reduction, however, was similar regardless of whether the ICD of Fat4 was present or not. These results suggested that actin polymerization is required for the maintenance of Fat4/Dchs1 boundaries, but that this requirement does not depend on the presence of Fat4 ICD.

**Figure 4:**
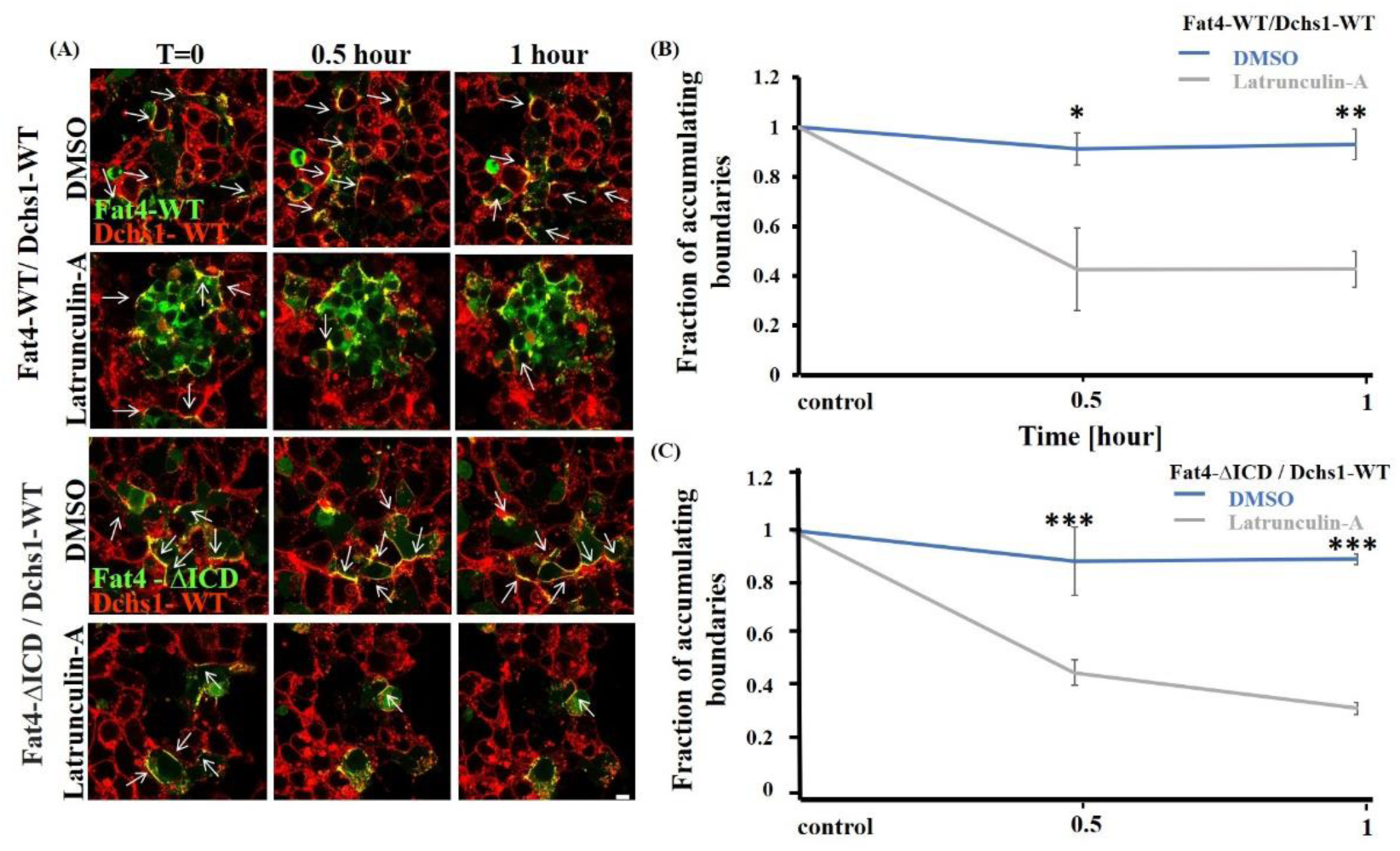
Actin depolymerization reduces the formation of Fat4/Dchs1 boundary complexes, but this reduction is independent on Fat4 ICD. **(A)** Images from co-culture experiments of Fat4-Citrine-WT (green): Dchs1-mCherry-WT (red) or Fat4-Citrine-ΔICD (green): Dchs1-mCherry-WT(red) Cells were treated with 1μM Lat-A or DMSO (control) for 0.5 or 1 hour. White arrows indicate boundary accumulation of Fat4/Dchs1 complexes. **(B-C)** Graphs showing the fraction of accumulating boundaries in co-culture of Fat4-Citrine-WT: Dchs1-mCherry-WT or Fat4-Citrine-ΔICD: Dchs1-mCherry-WT at different time points following treatment with Lat-A (gray) or DMSO (blue). All cultures were seeded for 24 hours before imaging. The fraction of accumulating boundaries was normalized to the average number of boundaries at t=0. Number of images: for the first-row n=20,18,16, for the second-row n= 18,16,18, for the third-row n=11,13,10, for the last row n= 15,14,13 respectively. Images taken from three independent experiments. Error bars represent S.E.M.*p< 0.05; ** P< 0.01, ***P<0. 001.Scale bars-10 μm.

### Actin accumulation is not correlated with Fat4/Dchs1 boundary accumulation

Since actin polymerization was required for the maintenance of the Fat4/Dchs1 boundary accumulation, we decided to test if the dynamics of actin cytoskeleton next to these boundaries is modified by the presence or absence of the Fat4 ICD. To do that, we introduced mCerulean3-LifeAct reporter, which labels actin in living cells, into our cells. We generated a construct which fused mCerulean3 to LifeAct, so that we can image actin, Fat4 and Dchs1 all at the same time. We then generated a stable cell line expressing both Fat4-Citrine-WT and mCerulean3-LifeAct, and another stable cell line expressing Fat4-Citrine-ΔICD and mCerulean3-LifeAct (Supplemental Fig. S4). We then looked for changes in the cytoskeleton dynamics in response to the presence or absence of Fat4 ICD in co-cultures of Fat4-Citrine-WT/mCerulean3-LifeAct: Dchs1-mCherry-WT (Fig. 5A-B) and Fat4-Citrine-ΔICD/mCerulean3-LifeAct: Dchs1-mCherry-WT (Fig. 5C-D). We tested whether the actin cytoskeleton reorganized around boundaries in different ways with or without Fat4-ICD, and whether it co-localizes with the Fat4 in the different co-cultures. Our live imaging movies showed that in some Fat4/Dchs1 boundaries we do observe actin co-localized next to the boundary (Fig. 5A, C’’’), while in other Fat4/Dchs1 boundaries we do not observe such actin co-localization (Fig. 5B, D’’’). We also did not see a clear difference between co-cultures that used WT Fat4 (Fig. 5A, B’’’) to those that used Fat4-ΔICD (Fig. 5C, D’’’). Overall, we did not find clear correlations between Fat4/Dchs1 accumulation and local actin dynamics, either in the presence or absence of Fat4 ICD.

**Figure 5:**
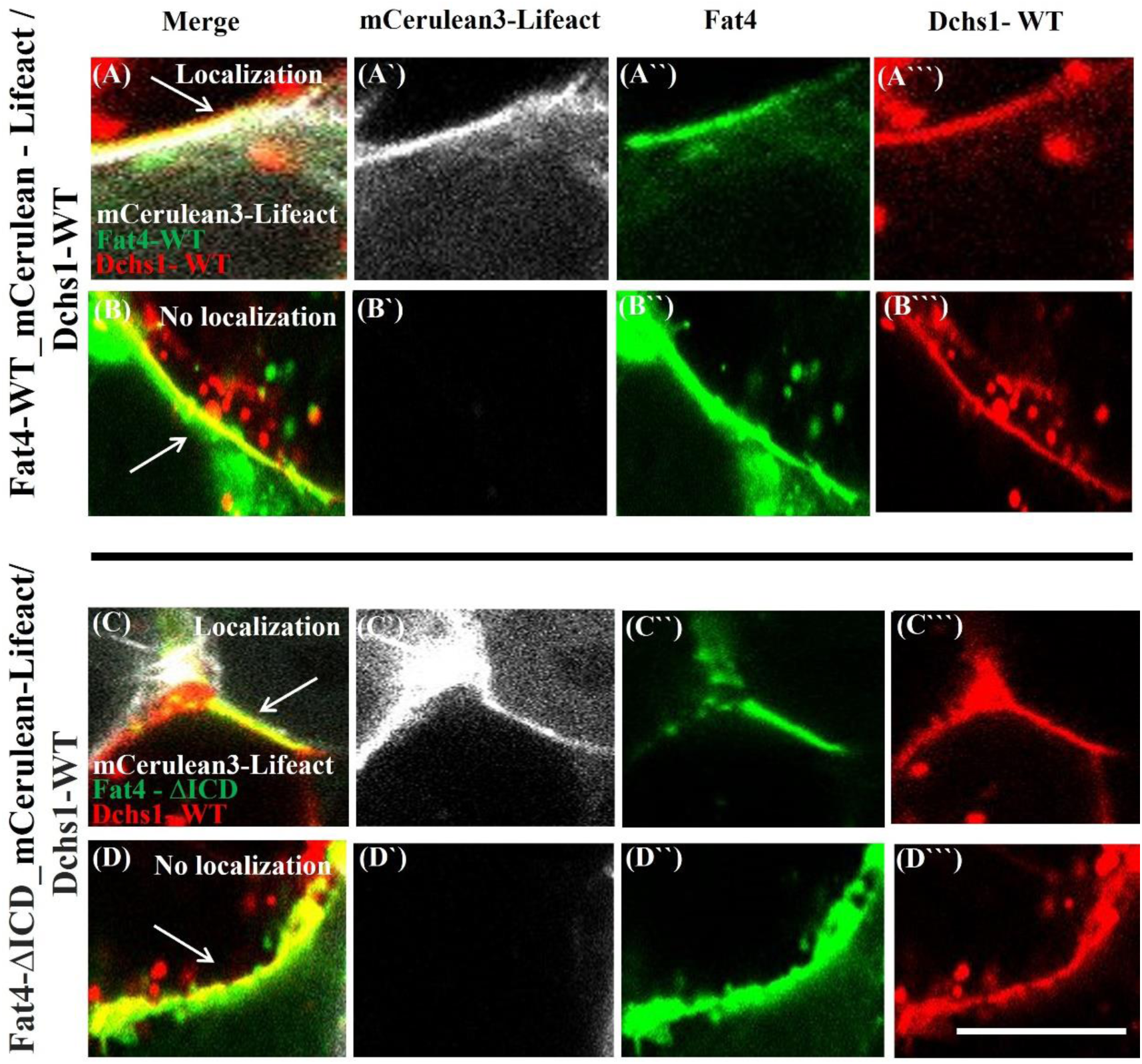
Actin dynamics are not affected by the presence or absence of the Fat4 ICD. **(A-D)** Images from a co-culture of Fat4-Citrine-WT(green)/mCerulean3-LifeAct(gray) or Fat4-Citrine-ΔICD (green)/mCerulean3-LifeAct(gray) cells with Dchs1-mCherry-WT (red). See a zoomed-out image in Fig. S4. White arrows represent localization of mCerulean3-LifeAct(gray) with Fat4/Dchs1 complexes. Scale bars-10 μm.

## Discussion

This work provides a mechanistic framework for understanding the roles of the ICD of Fat4 and Dchs1 in regulating the membrane dynamics of these complexes. Previous work in *Drosophila* has shown that Ft ICD is essential for both PCP and growth control and specific domains within the Ft ICD were associated with each function (13–15). On the other hand, the ICD of *Drosophila* Ds did not seem to be essential for PCP (13). Replacing the *Drosophila* Ft ICD with mammalian Fat4 ICD showed that the former is only associated with PCP and not in growth control(15). Our results show that the ICD of Fat4 does plays a role in trans-endocytosis and recycling of Fat4-Dchs1 complexes. However, the ICD of Dchs1 does not seem to affect either of these functions, consistent with the observations in *Drosophila*.

The observation that the ICD of Fat4 is required for TEC of boundary complexes, raises the question of what is the role of TEC in coordinating PCP. A Previous work on core pathway complexes (Fz6/Vangl2/ Celsr1) in the epidermis has shown that TEC is used for quickly remodeling cell boundaries during cell divisions (23). Previous works and the current work, in both *Drosophila* and mammalian cell culture, show that Ft/Ds and Fat4/Dchs1 boundary complexes are extremely stable (16,24–25). We therefore suggest that TEC in the Fat4/Dchs1 system may also be important for plasticity and remodeling of cell boundaries in different conditions, as it allows removing highly stable complexes in a very efficient manner. The identified role of the ICD of Fat4 in TEC suggests that the TEC is a regulated process controlled by some yet unknown interactors of the Fat4 ICD.

Consistent with the idea that Fat4 ICD is important for recycling of Fat4/Dchs1 complexes, we also find that the turnover of boundary complexes upon Ca**^+2^** depletion is faster with WT Fat4 than with Fat4 ΔICD.

The observations that WT Fat4 exhibit higher TEC and faster turnover seem to be somewhat at odds with the observation that Fat4/Dchs1 complexes accumulate more with WT Fat4 than with Fat4 ΔICD (since faster turnover is expected to lead to lower accumulation). This discrepancy may suggest that the ICD of Fat4 also regulate the complex formation and not only its degradation or turnover. However, we do not have direct evidence to support this conclusion.

A likely candidate for regulating trans-endocytosis and membrane complex turnover is the actin cytoskeleton. Previous work on Ephrin mediated trans-endocytosis, showed local enrichment of F-actin at EphB2 internalization sites (26). Our results show that blocking actin polymerization by Lat-A reduces Fat4/Dchs1 boundary accumulation, however, we do not see a difference between experiments using WT Fat4 and Fat4 ΔICD. Moreover, we do not identify specific actin localization or enrichment next to Fat4/Dchs1 boundaries. This suggests that interaction of actin with Fat4 ICD is not a crucial element in regulating TEC and Fat4/Dchs1 complex turnover.

Overall, our observations and conclusions provide insights into the dynamics underlying the formation and maintenance of Fat4/Dchs1 complexes at the cell boundary, and how it is controlled by the ICD of Fat4. Turnover dynamics of stable boundary complexes may be a crucial element for the establishment and plasticity of PCP. Such turnover may require exerting mechanical forces that lead to internalization of boundary complexes. Additional tools may be required for measuring and perturbing such forces. The insights from our work may also be relevant for other boundary complexes formed by cell adhesion proteins such as cadherins, nectins, and other protocadherins.

## Conclusions

In summary, this study provides a mechanistic framework for understanding the roles of the ICD of both Fat4 and Dchs1 in regulating the dynamics of Fat4/Dchs1 membrane complexes associated with planar cell polarity. The ICD of Fat4 is important for trans-endocytosis and turnover of membrane Fat4/Dch1 complexes, which may contribute to the plasticity of PCP under different conditions.

## Acknowledgements

We acknowledge the expert advice of Helen McNeill suggesting the Ca2+ turnover experiments and providing support.

## Author Contributions

This study was conceived and planned by Y.E. and D.S. Experiments were performed and analyzed by Y.E. Reagents and advice were provided by O.L. Code for analysis was developed by R. C. The manuscript was written by Y.E and D.S.

