## Supplemental information for "The Fat4 intracellular domain controls turnover of Fat4/Dchs1 planar polarity membrane complexes"

### Supplementary

| Cell line | Base cell line | Plasmid | Selection marker | Sequences |
|---|---|---|---|---|
| Fat4-Citrine-WT | HEK293T | Fat4-Citrine-WT | Zeocin | https://benchling.com/s/seq-eXJdY2B10zo8n1K6mJcu?m=slm-7dto4fVicfgDM0ChM27s |
| Fat4-Citrine-ΔICD | HEK293T | Dchs1-mCherry-WT | Zeocin | https://benchling.com/s/seq-1zPtzt19xUtxZsn1TYYu?m=slm-Nbs0ocADkFu4e058yw4T |
| Dchs1-mCherry-WT | HEK293T | Fat4-Citrine-ΔICD | Hygromycin | https://benchling.com/s/seq-aBtc5bv1RfeD1awafeNS?m=slm-5iOLtM3Ajo44ikoEb8GG |
| Dchs1-mCherry-ΔICD | HEK293T | Dchs1-mCherry-ΔICD | Hygromycin | https://benchling.com/s/seq-mo6kLufTTdvTM4nUqh1V?m=slm-g1jFNAD5P92DtZHJlAEA |
| Fat4-Citrine-WT/ mcerulean3-LifeAct | HEK293T | mcerulean3-LifeAct | Zeocin +Blasticidin | https://benchling.com/s/seq-hSwSxSBUtfKzHnDEOv3h?m=slm-dcQEpvGxUjgszpGiwrYw |
| Fat4-Citrine-ΔICD/ mcerulean3-LifeAct | HEK293T | mcerulean3-LifeAct | Zeocin+ Blasticidin | https://benchling.com/s/seq-hSwSxSBUtfKzHnDEOv3h?m=slm-dcQEpvGxUjgszpGiwrYw |

**Supplementary Table 1: List of Plasmids**

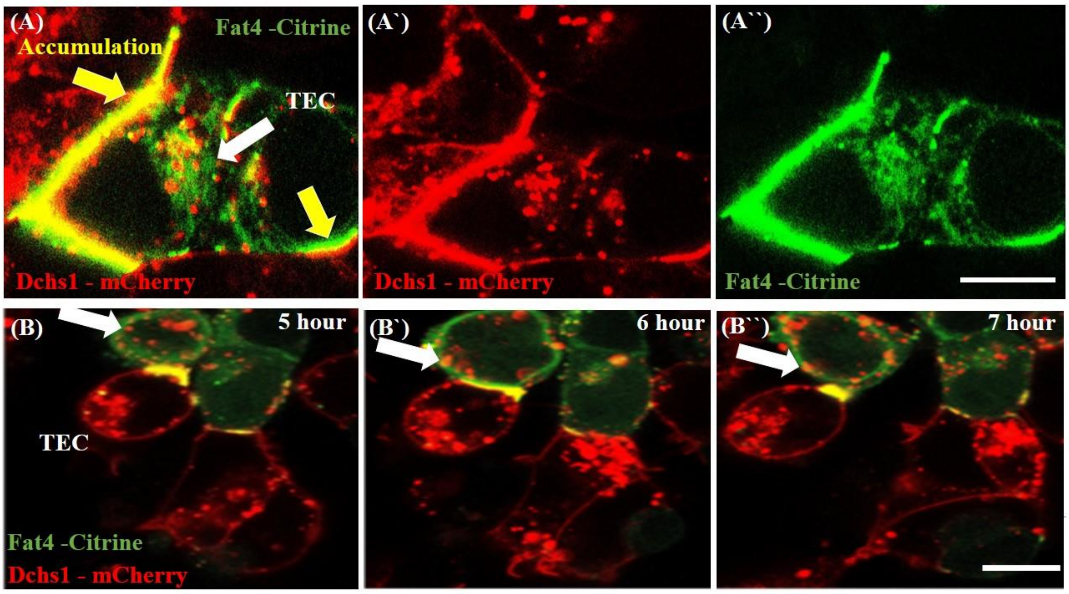


**Figure S1: Boundary accumulation and trans-endocytosis (TEC) is observed in co-cultures of Fat4-Citrine and Ds1-mCherry cells (A-A'')** A co-culture of Fat4-Citrine-WT (green): Dchs1-mCherry-WT (red) exhibits boundary accumulation (yellow arrows) and trans-endocytosis (TEC) (white arrow**). (B-B'')** A filmstrip showing the trans-endocytosed vesicles dynamically move inside Fat4-Citrine cells (white arrows). Time from beginning of the movie is indicated. Scale bars-10 μm

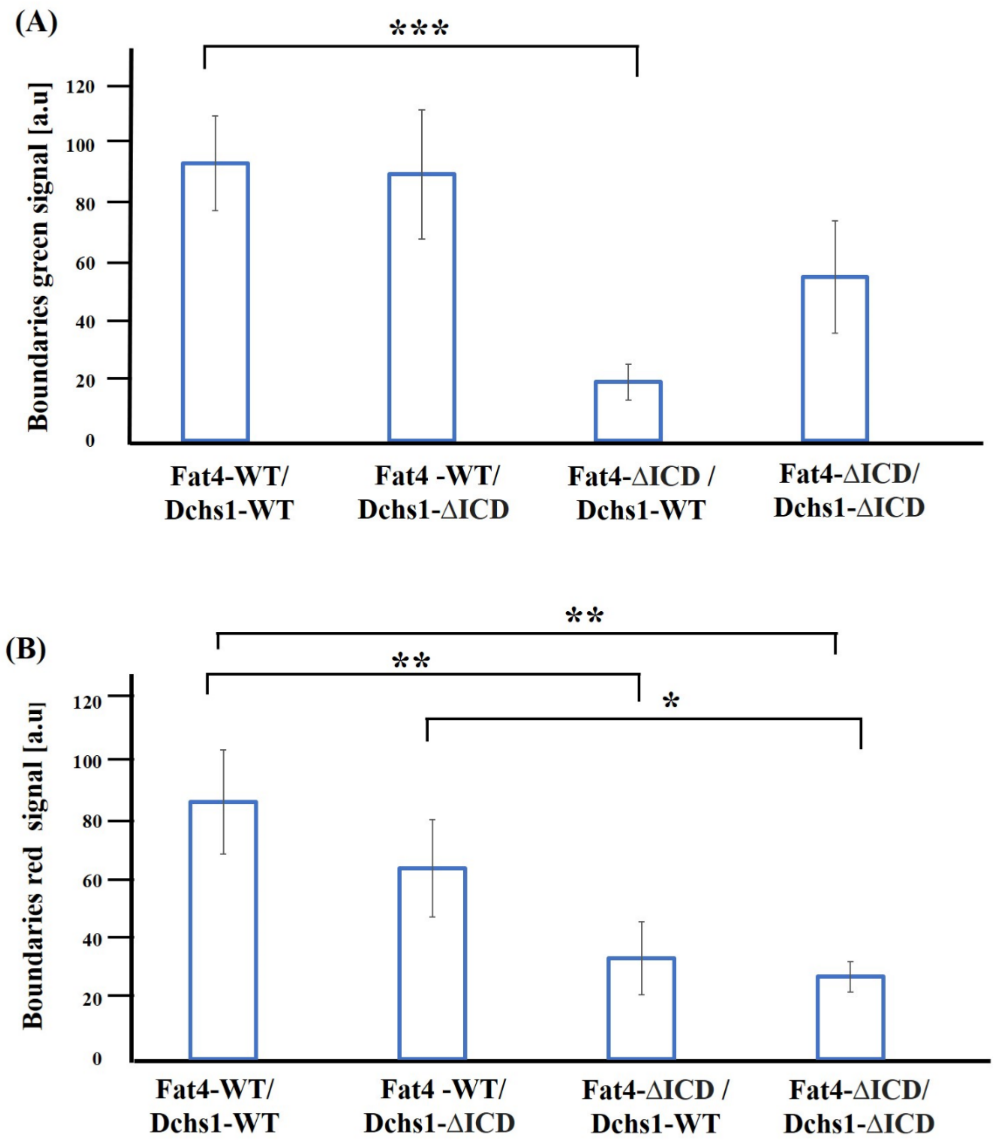


**Figure S2: The ICD of Fat4 affects accumulation of Fat4 and Dchs1 on the boundary. (A, B)** Measurement of the boundary Fat4-Citrine (green) and Dchs1-mCherry (red) in the different co-cultures (as indicated)**.** Values are the mean of three independent repeats. Number of images: for Fat4-Citrine-WT: Dchs1-mCherry-WT n=30, For Fat4-Citrine-ΔICD: Dchs1-mCherry-WT n=23, for Fat4-Citrine-WT:Dchs1-mCherry-ΔICD n=19, Fat4-Citrine-ΔICD: Dchs1-mCherry-ΔICD n=15. (P-values: * $< 0.05$; ** $< 0.01$, ***$<0.001$)

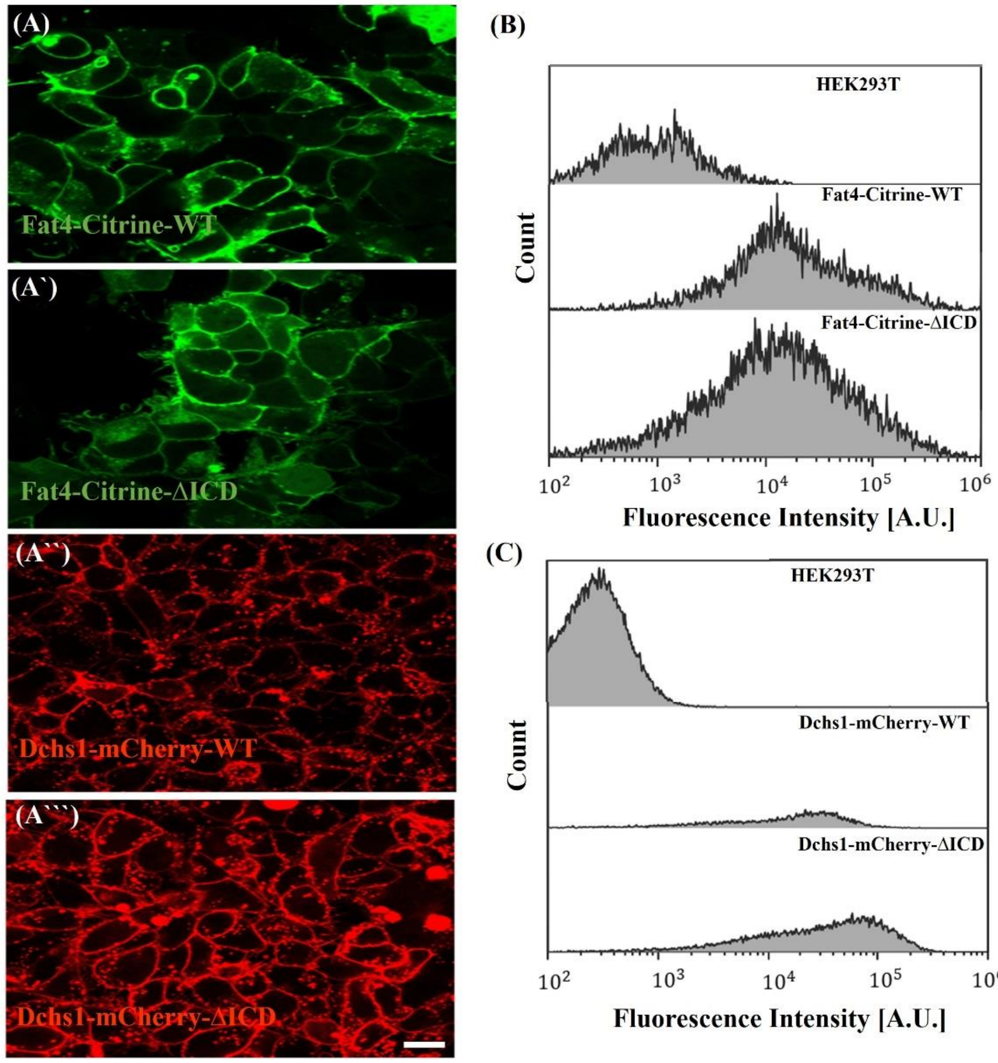


**Figure S3. Expression and localization of Fat4 and Dchs1 are not affected by absence of ICD (A-A''')** Representative images of Fat4-Citrine-WT, Fat4-Citrine-ΔICD, Dchs1-mCherry-WT, and Dchs1-mCherry-ΔICD shows similar subcellular localization of WT and ΔICD mutant variants**.** **(B)** Comparison of the fluorescence of Fat4-Citrine-WT and Fat4-Citrine-ΔICD. Control cells that do not express fluorescent protein (HEK293T cells) are shown in the top row. **(C)** Comparison of the fluorescence of Dchs1-mCherry-WT and Dchs1-mCherry-ΔICD cells. Control cells that do not express fluorescent proteins (HEK293T) are shown in the top row. Scale bars-10μm.

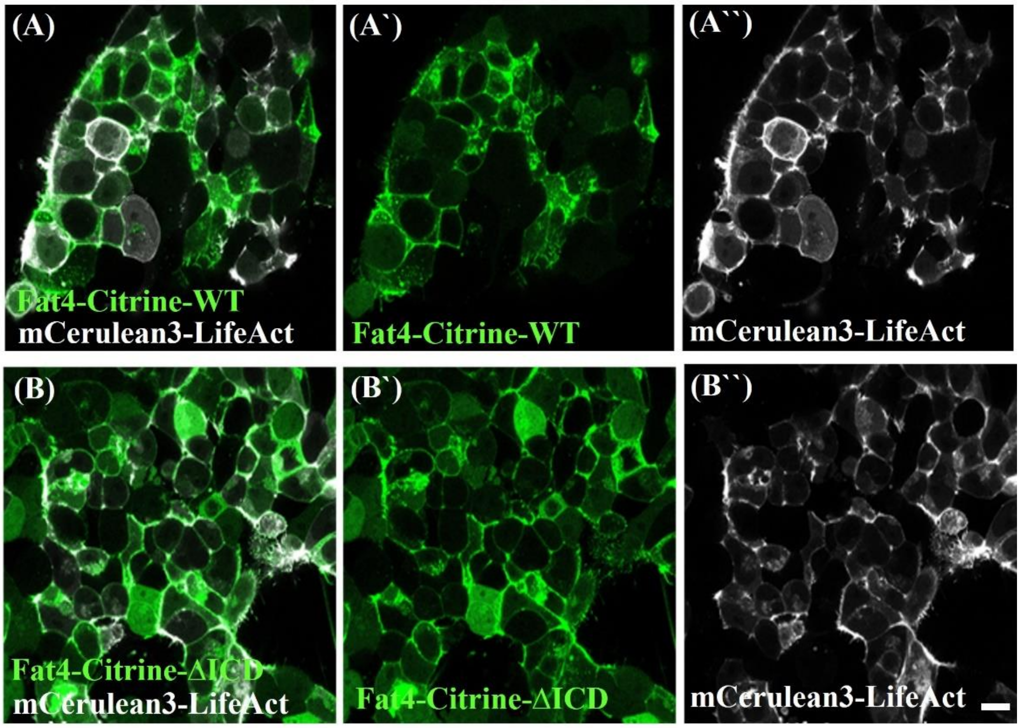


**Figure S4. Cell lines expressing both Fat4 variants and LifeAct reporter. (A)** Representative images of Fat4-Citrine-WT (green)/mCerulean3-LifeAct(gray). **(B)** Representative images of Fat4-Citrine-ΔICD (green)/ mCerulean3-LifeAct(gray). Scale bars-10 μm.

**Movie S1: Time-lapse FRAP movie** showing the dynamics of Fat4 in the complex of Fat4-WT/Dchs1-WT on the accumulating boundary.
Movie used to generate filmstrip in Figure 2A.

**Movie S2: Time-lapse FRAP movie** showing the dynamics of Fat4-ΔICD in the complex of Fat4-ΔICD /Dchs1-WT on the accumulating boundary.
Movie used to generate filmstrip in Figure 2D.
